# Predicting optimal pollinator visitation rates to crops using simple mechanistic models

**DOI:** 10.64898/2026.11.28.769547

**Authors:** Ignasi Bartomeus, Jake Bishop, Mike Garratt, Ellen Knight, Tom D. Breeze, Maxime Eeraerts

## Abstract

While the positive effect of pollinators on crop production has been clearly demonstrated, characterising the shape of the relationship between pollinator visitation rates and crop reproductive success remains a challenge. Defining the functional form of this relationship is of pivotal importance to understand when pollination is limiting, and to develop operational optimal visitation rate values that can be used by practitioners to rapidly assess expected pollination levels based on pollinator activity and react accordingly. Given the large diversity of crops and cultivars, conducting detailed experiments for each of them is unlikely to be achieved in the short term or to keep pace with cultivar development. Here, we propose and test a mechanistic model based on biological first principles to infer optimal pollinator visitation rates for multiple crops. First, we show that a non-inflected functional form fits empirical data well. Second, we develop and test a flexible model to produce these functional forms from easy-to-obtain plant and pollinator information such as plant selfing rates, ovule number, and pollinator single-visit efficiency. Finally, we apply our framework to 25 major pollinator-dependent cultivated crops and illustrate its flexibility by using blueberries as a case study to show the advantages and limitations of the framework. We provide an R package to facilitate the uptake of this modelling framework by practitioners.

## Introduction

Agronomic research has a long history of quantifying the inputs needed for optimizing crop production. For example, control trials have provided clear guidelines on optimal fertilizer use, irrigation, or pesticide application (Nakachew et al. 2024, Singh 2014, Leach et al. 2025). By contrast, insect pollination remained hard to measure and was traditionally assumed to be either non-limiting or easily covered by stocking managed pollinators *ad libitum* (Garibaldi et al, 2020). However, in the last 20 years, the importance of both managed and wild pollinators has been made evident, mainly through synthesis papers showing a positive relationship between pollinator visitation and crop production (Garibaldi et al. 2013, Rader et al. 2016, Reilly et al. 2024). Consequently, the interest in managing crop pollination with both managed and wild pollinators has increased (Garibaldi et al. 2014). Despite these efforts, the intrinsic variability across crops and cultivars (Reilly et al. 2024; Eeraerts et al. 2024), the assumption of linear relationships between pollinator visitation and yield effects (Eeraerts et al. 2025), and the confounding factors affecting yield beyond pollinators (Deguines et al. 2014, Garratt et al. 2018), have prevented the emergence of clear guidelines for supporting and maintaining optimal pollinator visitation rates for different crops (Garibaldi et al 2020). Failing to address insufficient pollinator visitation risks overlooking the implementation of management interventions until economically significant pollination deficits have already compromised the production of economically viable outputs (Garratt et al. 2021, Eeraerts et al. 2024).

Crop production is affected by many factors, including fertilizers, water availability, or cropping density (e.g. Pfuderer et al 2026). Hence, observational studies risk confounding the effects of pollination on crops when pollinator availability is correlated with any of these factors (Chabert et al. 2026, Aizen et al. 2023). Moreover, most analyses make statistical simplifications such as assuming a linear relationship between pollinator visitation and crop production, despite having been shown that a non-linear relationship is more biologically likely and often fits the data better (Morris et al 2010, Eeraerts et al. 2025). The consequence is that even simple metrics, such as how many visits per flower are needed to ensure pollination is not limiting, are hard to extract from this type of observational analysis, even when crops are analyzed individually (e.g. Chabert et al. 2022). As an alternative, controlled experiments, where crop resources are kept constant and the number of visits per flower is tracked, can provide detailed information on how many visits by different pollinators are needed to maximize crop yield (Garratt et al. 2014, Garratt et al. 2016, Kendall et al. 2022; Kandori et al. 2024; Di Trani et al. 2024). However, these experiments are very time-consuming, and hence have only been done for a handful of crops. Even within those controlled experiments, striking differences are observed depending on the cultivar, environmental conditions used, and the pollinator community available, indicating that the findings are unlikely to be generalisable. For example, Kendall et al. (2022) found that the number of visits required for 90% fruit set varied from three to five for northern highbush blueberry and southern highbush blueberry, respectively, whereas for rabbiteye blueberry a fruit set of only 58% was reached after 15 visits. Overall, this suggests that results might be hard to generalize.

Here, we propose a well-known functional form relating pollinator visitation to crop reproductive success: the non-inflected curve (Morris et al. 2010), which is a Michaelis-Menten functional type. The advantage of this functional form over statistical quadratic fits is that its parameters have biological meaning. The curve is defined by three parameters which govern the relationship between pollinator visitation rates and crop seed set. Parameter *a* sets the y-intercept (i.e., seed set for unvisited flowers), parameter *b* sets the asymptote (i.e., maximum seed set attainable by a single flower), and parameter *c* governs the rate of approach to the asymptote (Fig. 1). All these parameters can be estimated from experimental or observational data, but can also be derived from first principles. For example, existing databases on selfing rates can inform about the expected intercepts (Klein et al. 2007, Siopa et al. 2024). Similarly, the maximum seed set attainable per flower can be approximated using the number of fertile ovules, information often present in botanical or agronomical manuals, or databases on the maximum levels of pollination (Gazzea et al. 2023; Siopa et al. 2024). Finally, the parameter describing the rate of approach to the asymptote can be derived from pollinator efficiency on a per-visit basis (Page et al. 2021; Földesi et al. 2021). This opens the door to derive mechanistic relationships for a wide variety of crops and contexts.

**Figure 1:**
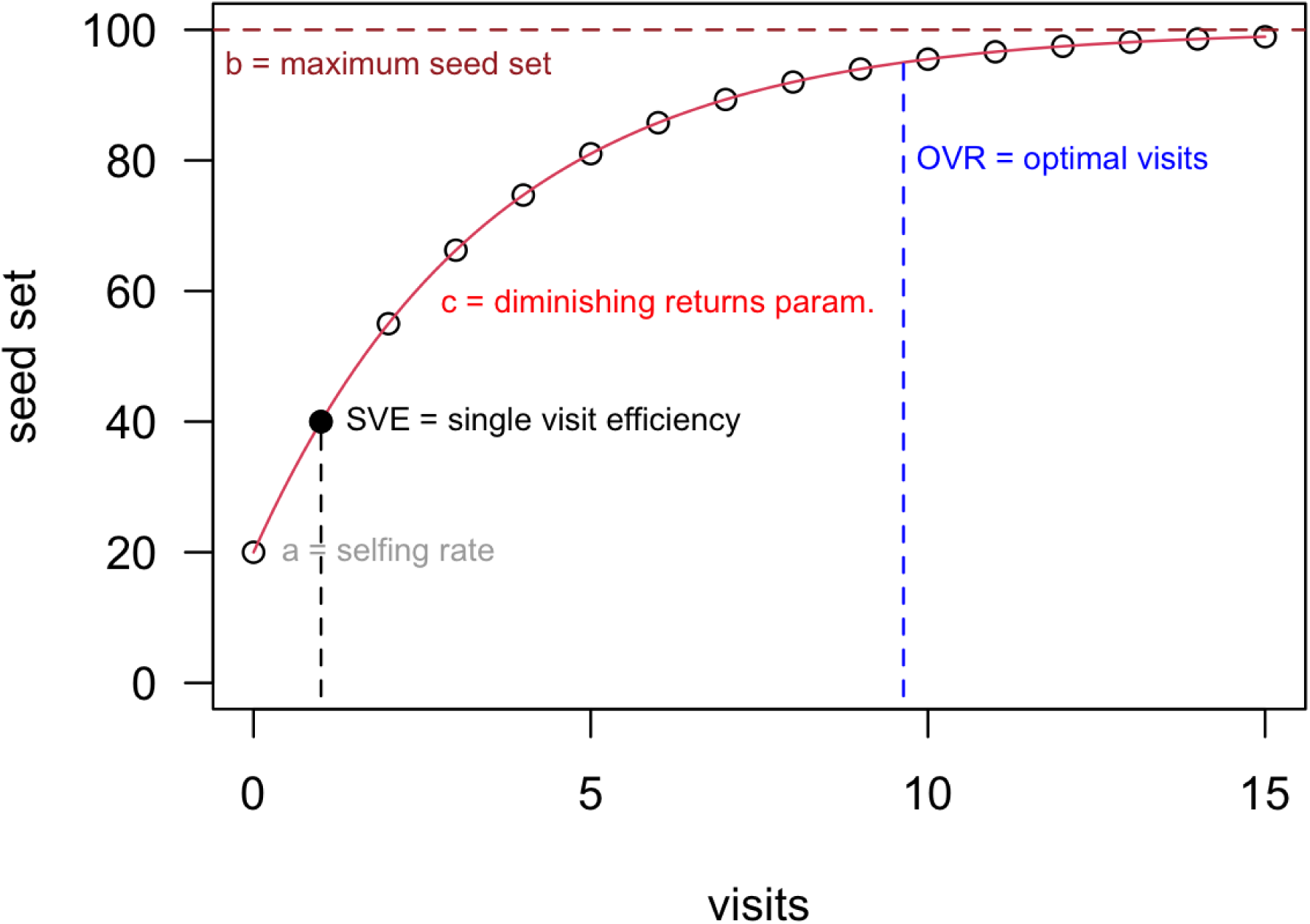
Theoretical explanation of the modelling framework for the relationship between pollinator visitation and seed set (%). The plot shows the intercept value defined as *a* (in grey), which is defined as the percentage of seeds obtained with no pollinator-mediated pollination (i.e. selfing). Parameter *b* defines the maximum attainable seed set (in brown). Parameter *c* describes the form of the curve (in red). We also show how single visit efficiency (*SVE*, in black) can be used to fit the curve, and the optimal visitation rate (*OVR*, in blue) is defined as the number of visits required to attain a given percentage of the maximum seed set (in the picture, the 95% is shown for illustrative purposes, but analysis of this paper use the 80%).

Defining a clear framework on how to assess pollinator contributions to a crops seed set will incentivise farmers and agronomists to assess pollination needs, underpin evidence based guidelines for pollinator management, and help identify areas where crop pollination is particularly strong or fragile. Here, we provide functions to calculate optimal visitation rates from experimental data when available, and from easy-to-obtain biological parameters when detailed experimental data is missing. By using a mechanistic model, we provide tools to rapidly assess pollination services to any given crop in any specific context. To facilitate the uptake of these tools, an R package accompanies this article, along with illustrative case studies showing its flexibility and limitations.

## Methods

### Modelling framework

We use a functional form proposed by Morris et al. (2010) that describes a seed set model that accounts for two different ways a plant can produce seeds: through autonomous self-pollination and through pollinator-assisted fertilization. In mathematical notation, the equation looks like this:

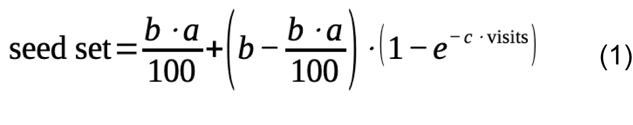

The formula is split into two main parts: a fixed baseline and a variable growth curve. The baseline (i.e., the intercept, representing autonomous seed set production) is defined by *b*, the maximum potential seed set, and *a*, the percentage of that maximum that can be achieved through self-pollination (e.g., if *a=20*, the plant can produce 20% of its seeds without insect pollination). The variable growth curve represents the additional seed production contributed by pollinator visitation, represented as *c,* and governs the exponential saturation curve. It starts at *a* and climbs toward *b* as the number of visits increases. A higher *c* means the plant reaches its maximum seed set with very few pollinator visits. This model shows that there is a diminishing return on subsequent pollinator visits due to the saturation of stigmas, or stagnation of growing pollen tubes (Rollin and Garibaldi 2019). Note that if *visits = 0*, the entire second half of the equation becomes 0, leaving only the baseline. As *visits* increase, the seed set grows quickly at first, then slows down as it approaches the maximum limit *b*.

Optimal visitation rate (*OVR*) can be calculated as the visits required to achieve 80% (or any other chosen threshold) of potential seed set. We chose 80% here for illustrative purposes, as many fruits rarely develop all their ovules into seeds, even at optimal pollination. However, this threshold should be selected according to the question under investigation and the characteristics of the target crop species.

### Data

Experiments relating seed set per flower to the number of visits received are scarce, but we were able to identify four illustrative studies covering different crops. These four studies cover different crop types: apple (Garratt et al. 2016), strawberry (Kandori et al. 2024), northern highbush blueberry, southern highbush blueberry, rabbiteye blueberry (Kendall et al. 2022), and watermelon (Di Trani et al. 2024). For blueberry, we converted fruit weight to seed number based upon the correlation between the two variables (averaged equations across cultivars: seeds = (fruit weight -0.09) / 0.06)), as these two parameters are highly correlated for this crop (Strik and Vance, 2019). Using these data, we fitted three different models and compared them by AIC and R2. The three models are i) a linear relationship, ii) a quadratic relationship, and iii) the non-inflected functional relationship described above. Models were run using the functions *lm()* for the linear and quadratic relationship, and *nls()* for the non-inflicted functional relationship in base R (R Core Team 2025).

### Mechanistic model

As these data are rare in the literature and hard to obtain experimentally, we derived a mechanistic version of our modelling framework that relies on first principles and does not require extensive experimental data. To that end, we set *b* as the maximum seed set a flower can attain. This can be extracted from the crop’s basic biology, as for many crops it will match the per-flower ovule production, a parameter often easy to obtain from the literature or easily counted in the field (the latter indicating achievable seedset in a particular context). Parameter *a*, given in % of seed set without pollinators, can be approximated knowing a plants’ selfing rate. That is, how many seeds the flower produces when pollinator-mediated pollination is prevented. This information is available in the literature (e.g., Klein et al. 2007, Siopa et al 2024) or can be calculated with basic low-effort experiments for a target crop or cultivar. The more challenging parameter to obtain is *c*. For this, we rely on pollinator per visit efficiency. Many studies measure seed set after a single visit of a pollinator. These values can be used to extrapolate how many visits per flower are required for fertilizing all its ovules. The average pollinator efficiency (SVE) can be calculated with a weighted mean across the available pollinator community, which is the representative metric, or calculated for different pollinator groups (see case study below for an example). These data are already available for many crops (Page et al. 2021; Földesi et al. 2021), and are expected to have lower variability across pollinators than pollinator visitation rates (Vazquez et al. 2005 and Kleijn et al. 2015). Obtaining these data from field observations requires only a moderate effort (Dafni et al. 2005). After this first visit, each subsequent visit is typically less efficient (Kendall et al. 2022), hence, we can penalize each consecutive visit by a diminishing returns function. This penalization can be informed biologically, as we expect these diminishing returns to approach *b*. Hence, we define the diminishing returns efficiency function as:

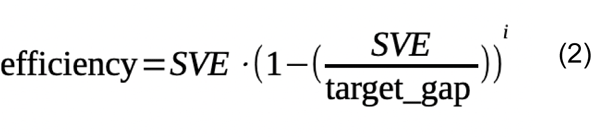

where the *target_gap* is the difference between the baseline and the maximum attainable seed set, and *i* is the number of extra seeds achieved after a single first visit. When *i = 0* (first visit), efficiency is equal to the single visit efficiency (SVE), but this value is exponentially lower at each subsequent visit.

To test the flexibility of this approach, we first compiled information on *a*, *b,* and *SVE* parameters for 25 major crops by scanning the literature (Table S1). Note that these are informative generic values, but we encourage users to look up or measure the values for their crop, cultivar, and region. Next, we compare these data for the four crops for which we have detailed data. Finally, we delve into the flexibility of the approach in a case study on blueberry pollination (Kendall et al. 2022). This allows us to demonstrate how values for different cultivars or pollinator communities can be customised.

## Results

The non-inflected functional form shows better or equal performance than the quadratic or linear relationship for all four crops (Fig. 2), demonstrating that it is a useful modelling framework. Despite needing to estimate three parameters (*a*, *b*, and *c*) against the two parameters of a linear model, AIC values are lower for the non-inflected fit for apple and blueberry (ΔAIC > 10) and similar for strawberry and watermelon (ΔAIC ∼ 0; Table S2). R2 values were always higher for the non-inflected fit (Table S2). Most importantly, when visually inspecting the fitted lines, our model avoids the unrealistic linear increase of seed set beyond the maximum number of ovules (*b*), and the reduction of seed set at high visitation rates shown by some quadratic fits. While such a seed set reduction has been shown in some crops (e.g. Saez et al. 2014), the mechanisms are often different (e.g., stigma damage) and the extent of these effects remain largely underexplored for most crops. In any case, these potential negative effects of higher-than-needed visitation rates can be assessed by flagging visitation rate values that are well beyond the calculated optimum.

**Figure 2:**
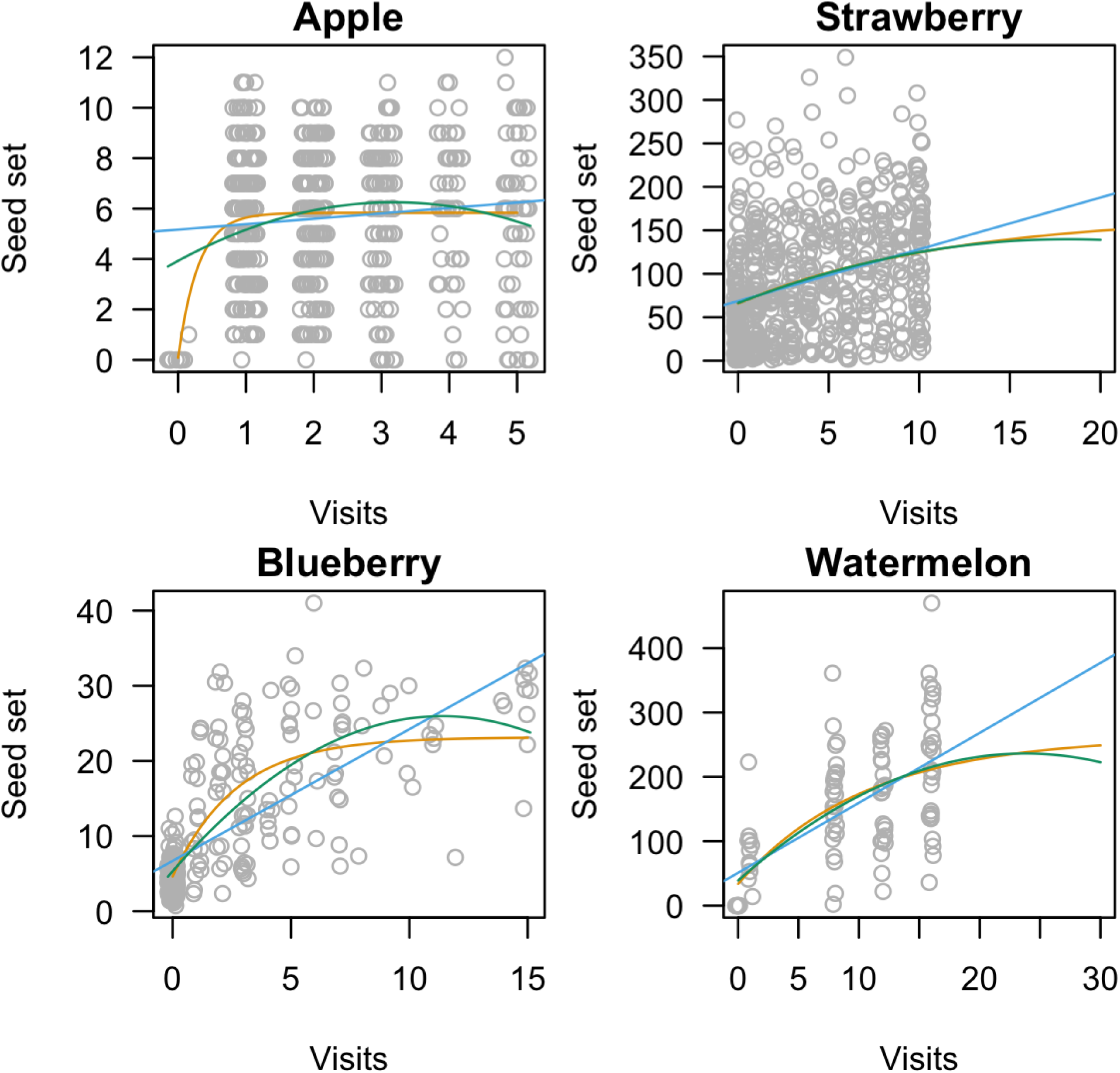
Seed set as a function of visitation rate for data collected on four different crops. a) apple, b) strawberry, c) southern highbush blueberry, and d) watermelon. Blue lines represent the linear fit, green lines the quadratic fits, and orange lines the non-inflected functional form.

Large variability in the underlying data is observable in the associated uncertainty found in all models and crops, indicating that even in controlled experiments, there is a wide uncertainty in the effects of pollinator visitation. In particular, for watermelon and strawberry, we do not observe a clear saturation at the maximum number of visits recorded, indicating that higher visitation numbers should be measured to ensure the upper part of the curve is well covered. Note that if we mis-estimate the *b* parameter, we will also mis-estimate the optimal visitation rates value. For example, in watermelon, the open treatment averages 264 seeds, indicating saturation at values higher than 15 visits, but we cannot map this value to a certain number of visits.

Using the generalized modelling approach, we calculated functional forms (Fig. 3) and optimal visitation rates (*OVR*; Fig. 4) for 25 crops using the three parameters extracted from the literature (*a*, *b*, and *SVE; Table S1*). Optimal visitation rates range from more than 10 visits for blueberry, kiwi, peppers, or squash, to less than five visits for pears or apples. Note that *OVR* depends on the interplay of *a*, *b*, and *SVE*, and none of those units alone can predict its values well.

**Fig 3.**
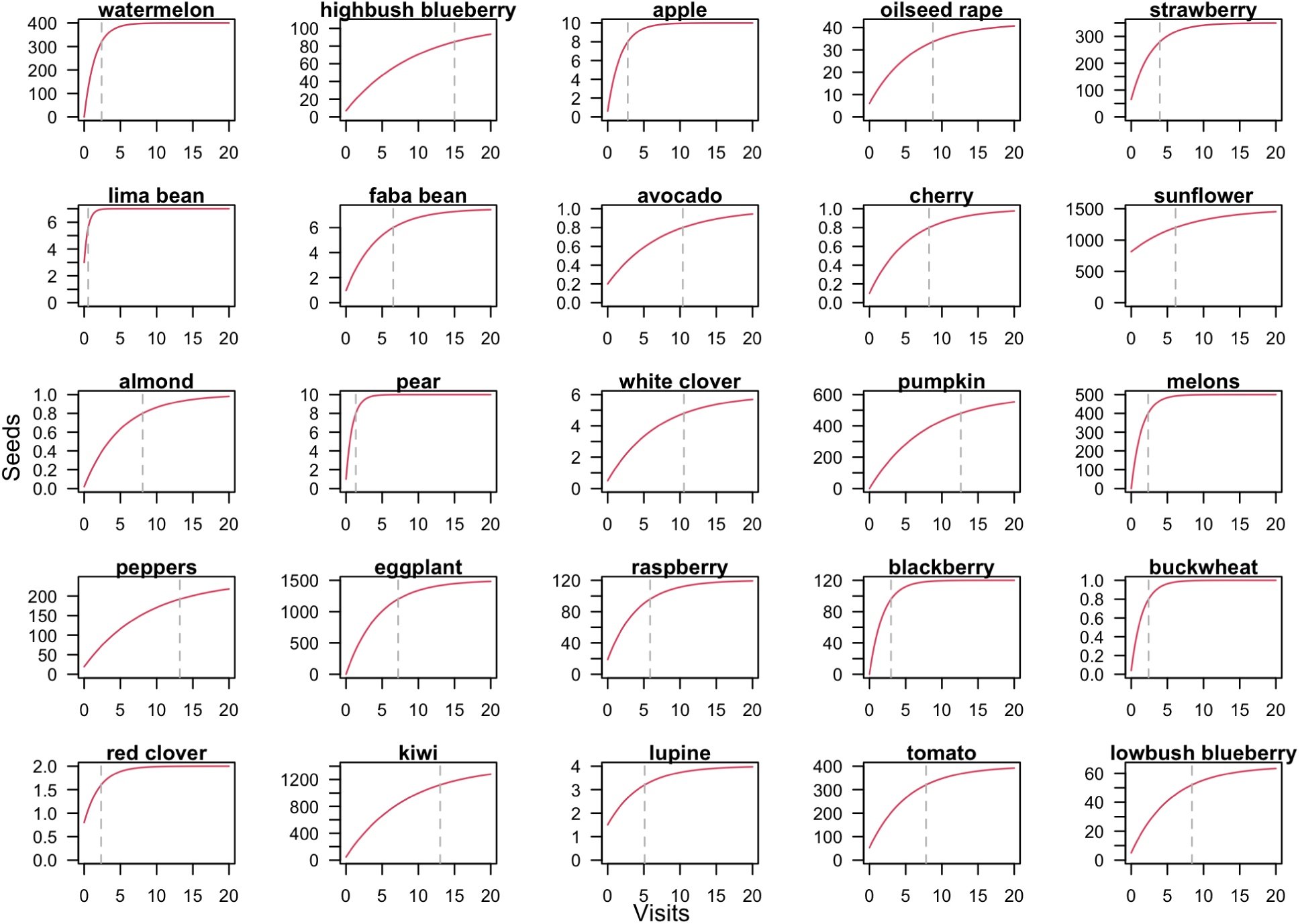
Derived functional forms for 25 major crops using a mechanistic model informed by biological parameters (Table S1). Grey dashed lines depict the optimal visitation rates to attain 80% of seed set.

**Figure 4:**
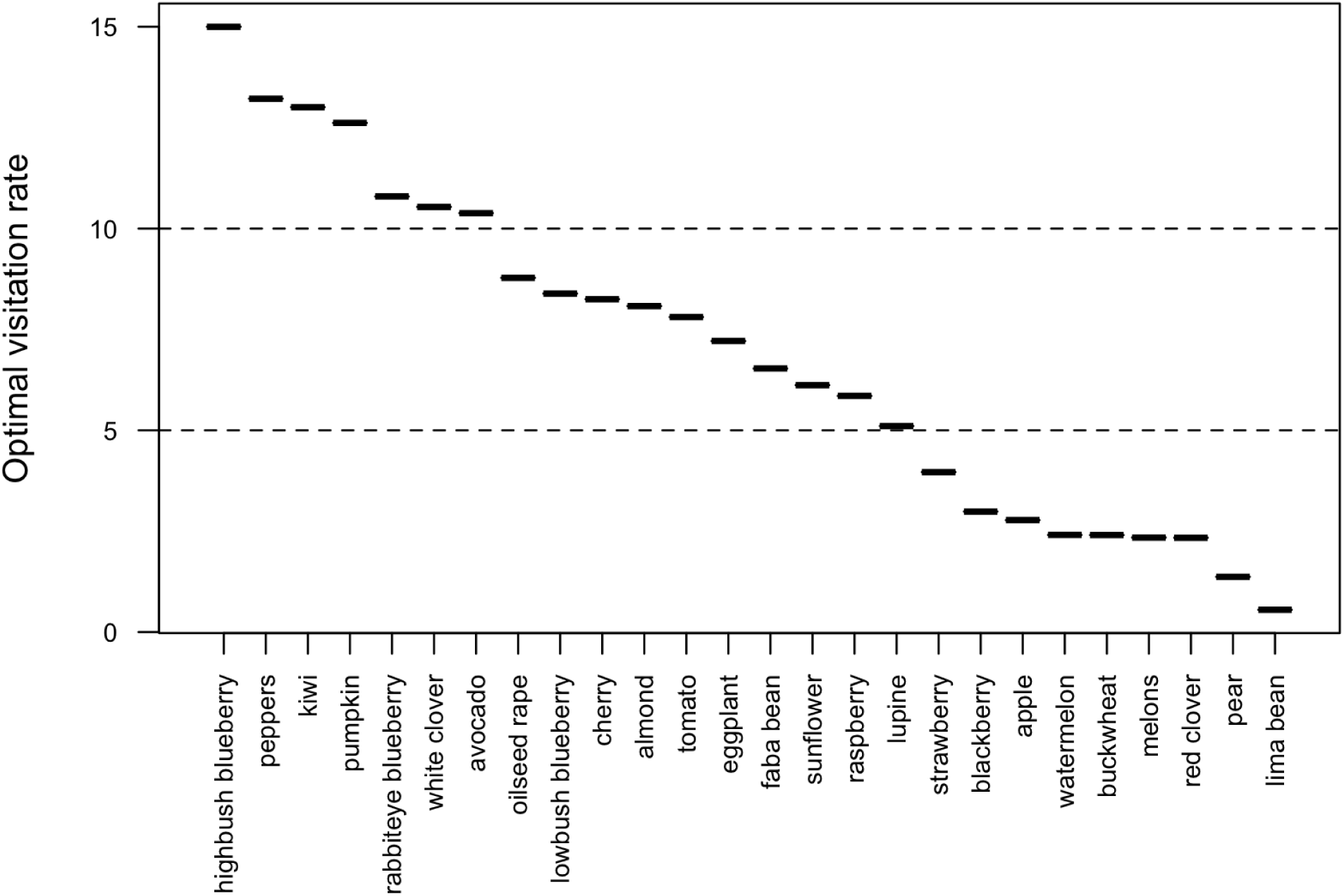
Model predictions of optimal visitation rates (*OVR*) for 25 main EU crops. *OVRs* are calculated to attain 80% of maximum pollination.

Focusing on the four crops for which we have sufficient empirical data, we can visually compare the model parameters for the data-driven values and mechanistic values. In general, there is low agreement (Fig. S1), as data from the data-driven and the empirical data come from different cultivars and/or contexts with different values. However, when we compare data-driven and mechanistic models for the same cultivars, the agreement is high (Fig. 5). *a* and *b* parameters are in general well estimated, but surprisingly, the selfing rates of the watermelon model are closer to the real value than the empirical fit (note that the empirical intercept does not capture the average of zero seeds in non-visited plants; Fig 2). On the other hand, *OVR* for watermelon tends to be lower for the mechanistic model than in the empirical data. This is because single-visit values in the data suggest > 70 seeds for a single visit, quickly saturating the mechanistic response, while the empirical data lacks a clear asymptote due to a cap on the number of visits measured, suggesting more visits are needed (up to 15). This highlights the importance of choosing the initial parameters well for mechanistic models, but also that data-driven models are not necessarily more robust due to the large variability in the data and the difficulty covering all the visitation range (e.g. for watermelon and strawberry).

**Figure 5:**
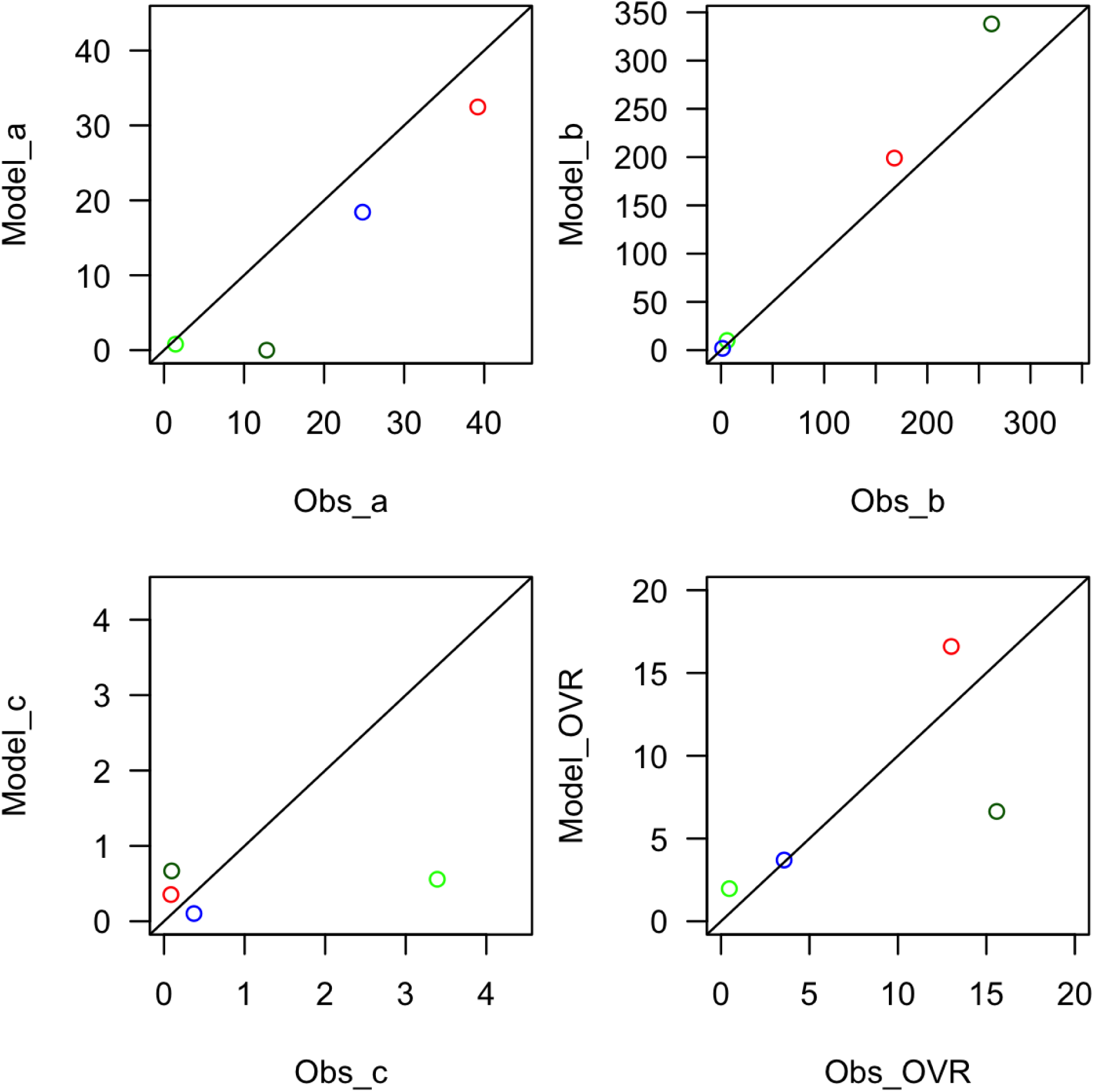
Comparison for the four main parameters (*a*, *b*, *c*, and *OVR*) for data-driven (Obs_ ; x axis) and mechanistic models (Model_ ; y axis). The 1:1 line is plotted. Light green circles represent apple, dark green represents watermelon, red represents strawberry and blue represents blueberry.

We show that with basic data, we can generate plausible functional forms for many crops and estimate their expected optimal visitation rates. However, as expected, these are at quite coarse resolution and should be treated as first approximations. Only 12 crops within the database had data on selfing rates and single-visit efficiency from the same study and cultivar (Table S1), while for other crops we only found this information for different cultivars or averages across cultivars. Given the large differences in pollination requirements of different cultivars (Siopa et al. 2024), it is risky to use generic values from different studies (Bishop and Nakagawa, 2020; Garratt et al. 2021; Siopa et al. 2024; Eeraerts et al. 2024). For example, when using Siopa et al. (2024) pollinator dependence averages per crop, and matching those with single-visit efficiency studies conducted in a single cultivar, we found 11 cases where seed set was greater for flowers with no visits than after a single visit, which makes little biological sense and likely reflects within crop variability. Hence, researcher experience and knowledge of the specific cultivar will drastically improve predictions.

Our framework allows for refinement of these predictions if appropriate data are available. Using detailed data collected by Kendall et al. (2022) for blueberry, we first predict the relationship between visitation and seed set for different blueberry crops (Fig. 6A), and second for different pollinator communities for southern highbush blueberry (Fig. 6B). We see that detailed information on selfing rates and maximum yield changes the predictions for the different blueberry crops. Similarly, different pollinator communities might affect the number of visits needed to maximise fruit production. Specifically, for southern highbush blueberry, receiving only honeybee visits requires more visits than receiving only *Tetragonula carbonaria* visits, with mixed communities showing an intermediate result.

**Figure 6:**
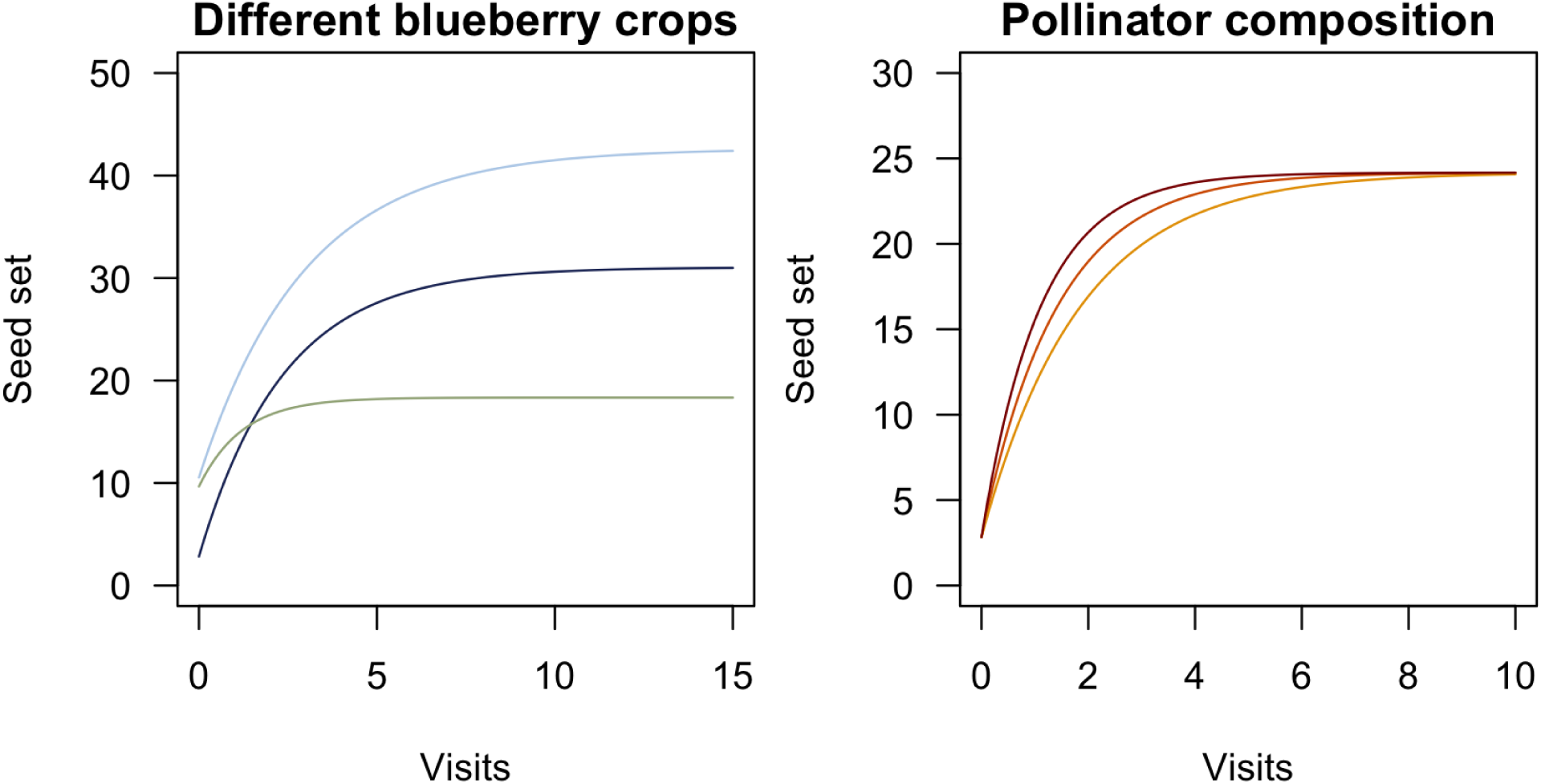
We use blueberry as a case study to illustrate how to go from generic predictions to tailored predictions to your context. Using blueberry data from Kendall et al 2022, we show A) that we can predict curves for different crops (green: northern highbush; dark blue: southern highbush; and pale blue: rabbiteye), and that B) different pollinator communities (yellow: honeybee-dominated; brown: wild bee-dominated; and red: mixed) in southern highbush blueberry show different curves.

## Discussion

We show how modelling the relationship between pollinator flower visitation rates and seed production is not only possible, but efficient and scalable. By developing a simple mechanistic model based on ecological first principles, we present generic optimal pollinator values for 25 common crops, and show how these values can be fine-tuned to specific cultivars and contexts. This is especially important given the high variability expected for these relationships, which depend on the cultivar of interest, the available pollinator community, and other contextual factors (Tamburini et al. 2019).

A core assumption of our modelling framework is that only pollinator-mediated effects are modeled. Rather than a limitation, we see this as a feature of the model, given it is interpreted correctly. When assessing pollinator effects on yield (Reilly et al. 2024; Garratt et al. 2021; Eeraerts et al. 2025), our mechanistic model does not confound pollinator effects with variables such as fertilizers, water availability, or cropping density, which can mask effects on final yield (Marini et al. 2015; Tamburini et al. 2017; Eeraerts et al. Submitted; Pfuderer et al. 2026). Hence, rather than being directly comparable, observational data and our model outputs can complement each other to help discover if something else is limiting yield besides pollinator levels. Similarly, when developing recommendations for optimal pollinator levels (Garibaldi et al. 2020), the model assumes those are independent of other management practices, which are assumed to be not limiting by the model. Note that when relating visitation to seed set using observational data directly, confounding factors can still apply, as evidenced by the large variability observed in empirical data. In fact, this is precisely why linking visitation rates to crop reproductive success has been elusive to date, and even well-researched crops lack clear guidelines on what the optimal pollinator levels are.

Despite the clear framework we provide, the choice of the input parameters still requires careful consideration. Ideally, selfing rates, maximum attainable seed set, and pollinator efficiency should be collected for your cultivar and pollinator community of interest. When this is not possible, informed choices can provide good proxies, but given the large variability even within the same cultivar of a crop, caution is needed when selecting those parameters from literature or assuming similarities between related crops or varieties. An illustrative example is the particular high optimal visitation value (15 visits) obtained for highbush blueberry (Fig. 3), as it is in general a crop that should be very easy to pollinate given its high selfing rates (DeVetter et al. 2022). However, given that we extracted selfing rates (*a*) and *SVD* from a study using an old cultivar (O’Neal) which is probably more difficult to pollinate, and maximum attainable seeds (*b*) from the cultivar Bluecrop, a more recent cultivar, with higher seed numbers, we might be overestimating the real visitation rate needed. Something worth highlighting is that there is great variation in the single-visit efficiency parameter depending on the pollinator species for which it was measured (Roquer-Beni et al. 2022, Cortés-Rivas et al. 2023). As an example, we found that our model matches the empirical data expectations when we use information on selfing rates, maximum seed production, and pollinator single-visit efficiency from the same study, but not when using available single-visit efficiency data from other studies (Compare Fig. 5 and Fig. S1). This was also the case for cherry, where single visit efficiency was calculated for honeybees, bumblebees, solitary bees and hoverflies (Eeraerts et al. 2020), but hoverflies, honeybees and bumblebees had very low efficiency compared to solitary bees, increasing the number of total visits needed. Hence, the values provided for the 25 crops analyzed are generic values for these crops, but we expect them to vary across cultivars and pollinator communities within crops. In fact, our case study using blueberry data confirms how fine-tuning the input parameters to specific contexts can change the outputs in meaningful ways.

While we focus on the pathway between visitation rate per flower and seed set per fruit, we argue that expanding the framework to cover the full chain from pollinator abundance to yield is easily achievable. Most ecological datasets (Allen-Perkins et al. 2022) measure pollinator activity as visits per square meter and hour rather than in visits per flower. This is because the most common method is walking a fixed-length transect over a fixed time while surveying insects interacting with the flowers. Knowing the expected flowers per square meter for your field, flower longevity, and using simple conversion rates, we can easily convert visits per square meter per hour to an estimate of visits per flower (implemented in our R package as *V_transect_to_flower()*). Similarly, while we work at the flower level, we should think of this flower as an average flower. Visitation rates of less than one visit per flower, such as in almond, cherry, avocado or kiwi, are hence relevant, as those imply not all flowers will get at least one visit (e.g. Garratt et al. 2014 show not all oilseed rape flowers are visited). When scaling up from individual flowers to whole plants, as is implicit in estimating curves for single-seed fruits, it becomes particularly important to use maximum seed set values measured in the target crop (rather than e.g. a botanical definition) as across-flower seed set relates closely to yield and the contextual factors (e.g. agronomic factors) that constrain it. A second important step is to turn our seed set per flower into yield, for which we would (simply) need to know the relationship between seed set and both the number of fruits and the fruit weight. This can be, again, obtained from the literature (Strik and Vance 2019; Garratt et al. 2021), or calculated empirically using a few selected fruits covering a large range of weights for which we also know seed number. If desired, this relationship can be truncated to account for abortions if the data suggest fruits with fewer than a minimum number of seeds tend to abort. Then, calculating yield implies a simple conversion factor chain where: Yield = Flowers/Ha * Visits/Flowers * SeedSet/Visits * FruitWeight/SeedSet = Weight/Ha. Of course, the more parameters we estimate, the more uncertain the final value obtained becomes, and proper error propagation is needed (i.e., accounting for the error or variability in all parameter inputs). Other management practices like thinning might also complicate extrapolating to yield.

While we still lack high-quality data on the required input parameters (*a*, *b* and *SVE*) for most cultivars, we hope that having a unified framework will speed up the collection of this data and its addition to unified databases (e.g., CropPol (Allen-Perkins et al. 2022), PolLimCrop (Siopa et al. 2023)). In this paper, we provide the first raw estimates of how many visits per flower we expect different crops to need to ensure pollen is not limiting. This value is critical for starting to measure pollination at the same level as other agricultural inputs such as fertilizers or pesticides, and improve management beyond actual guesstimates (Garibaldi et al. 2020). With these guidelines, visitation rates well below the optimal recommendations will require measures to improve pollination services such as adding honeybees or other managed pollinators (Mallinger et al. 2021, Osterman et al. 2023), or enhancing on-field (e.g., flower strips; Albrecht et al. 2020) or off-field (e.g., landscape linear elements; Garratt et al. 2017; Garibaldi et al. 2014) habitats. On the contrary, values well above the optional recommendations indicate that managed honeybees might be at too high densities. Given the known negative effects of managed pollinators at high densities (Saéz et al 2014; Chirilá et al. 2026), this can be a powerful tool to convince farmers to reduce the numbers of deployed hives when necessary. In fact, some crops suffer negative effects when managed pollinators are too abundant, as stigmas can be damaged (Sáez et al. 2014) or the managed honeybees may displace more effective wild pollinators (Li et al. 2025). Finally, here we consider only the agronomic aspects of yield, further work can link our framework to assess the full economic impacts (including the effects on management costs) of passing above or below these optimal thresholds. On a per flower basis, sub-optimal pollination may result in monetary losses to growers, particularly for crops where minimum size and shape thresholds are required before the crop can be sold (e.g. strawberries) (Wietzke et al. 2018; Gazzea et al. 2023). On the other hand, upscaled to whole plants, excessive pollination may result in higher management costs such as increased thinning of apples to maintain quality or prevent branch damage (Garratt et al., 2014). Thus, what is optimal for the plant may not be optimal for growers.

We encourage researchers to use our modelling framework, for which a dedicated R package is provided. Only by contrasting modeled and observed data can we enhance models and make them useful (Dietze et al. 2018). It’s been more than five years since Garibaldi et al. (2020) proposed better monitoring of pollinator inputs in agro-ecosystems. While this paper has received more than 100 citations, just a handful of those actually provide critical thresholds for pollinator optimal visitation rates. The time is ripe for developing these numbers and using them in applied settings.

## Data availability

An R package with all functions is available here: https://github.com/ibartomeus/VisitationR (and deposited in Zenodo with DOI: https://zenodo.org/records/24158929). Code and Data to reproduce this paper is available here: https://github.com/ibartomeus/VisitationRates (and deposited in Zenodo: https://doi.org/10.5281/zenodo.24499316).

## Acknowledgements

We acknowledge funding from Horizon Europe Research and Innovation Programme projects AGRI4POL (grant agreement No 101181146) and VALOR (grant agreement: 101181169). IB thanks University of Reading for hosting him during September 2025.

## Authors’ contribution

Conceptualization: IB, JB, MG, ME

Data curation: ME, MG, IB

Formal analysis: IB

Writing – initial draft: IB, ME

Writing – Review and editing: ME, JB, MG, EK, TB

## Supplementary material

**Table S1:** Parameters extracted from the literature. *a* is in seed set, *b* in number of ovules, and SVE in seed set after a single visit of honeybees (HB), Bumblebees (BB), other wild bees (WB), hoverflies (HF), or other.

| **Crop** | **Scientific name** | **Cultivar** | **a** | **b** | **SVE_HB** | **SVE_BB** | **SVE_WB** | **SVE_HF** | **SVE_Other** |
|---|---|---|---|---|---|---|---|---|---|
| watermelon | *Citrullus lanatus* | Royal Jubilee | 0<sup>1</sup> | 400<sup>18</sup> | 160<sup>1</sup> | 230<sup>1</sup> | NA | NA | NA |
| highbush blueberry | *Vaccinium corymbosum* | O'Neal | 7<sup>2</sup> | 106<sup>19</sup> | 12<sup>2</sup> | 20<sup>2</sup> | 18<sup>2</sup> | NA | NA |
| apple | *Malus domestica* | Breaburn and Brampley | 0.6<sup>3</sup> | 10* | 5.8<sup>3</sup> | 5.8<sup>3</sup> | 6.1<sup>3</sup> | 0.8<sup>3</sup> | NA |
| oilseed rape | *Brassica napus* | Hyola 61 | 6<sup>4</sup> | 42<sup>20</sup> | 12<sup>4</sup> | NA | 11<sup>4</sup> | NA | NA |
| strawberry | *Fragaria × ananassa* | Tyonoka | 65<sup>5</sup> | 350<sup>21</sup> | 78<sup>5</sup> | NA | 53<sup>5</sup> | 90<sup>5</sup> | 79<sup>5</sup> |
| lima bean | *Phaseolus lunatus* | Multiple | 3* | 7<sup>22</sup> | NA | NA | 6.4<sup>29</sup> | NA | NA |
| faba bean | *Vicia faba* | Tiffany | 0.97<sup>6</sup> | 7.5* | 1.8<sup>6</sup> | 1.66<sup>6</sup> | 3.4<sup>6</sup> | NA | NA |
| avocado | *Persea americana* | Multiple | 0.2<sup>7</sup> | 1* | NA | NA | NA | NA | 0.3*<sup>,7</sup> |
| cherry | *Prunus avium* | Kordia, Regina | 0.1<sup>8</sup> | 1<sup>23</sup> | 0.1<sup>26</sup> | 0<sup>26</sup> | 0.25<sup>26</sup> | 0.1<sup>26</sup> | NA |
| sunflower | *Helianthus annuus* | Multiple | 813<sup>9</sup> | 1500* | 800* | 900* | 1000* | NA | NA |
| almond | *Prunus dulcis* | Multiple | 0.02<sup>10</sup> | 1* | 0.14<sup>27</sup> | NA | 0.25<sup>27</sup> | NA | NA |
| pear | *Pyrus communis* | Multiple | 1* | 10* | NA | NA | 7* | NA | NA |
| white clover | *Trifolium repens* | Multiple | 0.5* | 6<sup>24</sup> | 1.24<sup>24</sup> | NA | NA | NA | NA |
| pumpkin | *Cucurbita pepo* | Mystic Plus | 0<sup>11</sup> | 600* | 63.65<sup>11</sup> | 113<sup>11</sup> | 38.9<sup>11</sup> | NA | NA |
| melons | *Cucumis melo* | Multiple | 0 * | 500* | 222.8<sup>28</sup> | NA | 274<sup>28</sup> | NA | NA |
| peppers | *Capsicum annuum* | Maya | 19.3<sup>12</sup> | 240* | 35.2<sup>12</sup> | NA | 51.5<sup>12</sup> | NA | NA |
| eggplant | *Solanum melongena* | Multiple | 0* | 1500* | NA | NA | NA | NA | 300* |
| raspberry | *Rubus idaeus* | Cowichan, Latham, Royalty | 18.8<sup>13</sup> | 120<sup>13</sup> | 41.15<sup>13</sup> | 41<sup>13</sup> | 40.4<sup>13</sup> | NA | NA |
| blackberry | *Rubus fruticosus* | Multiple | 0* | 120* | NA | NA | NA | NA | 50* |
| buckwheat | *Fagopyrum esculentum* | Multiple | 0.04<sup>14</sup> | 1* | NA | NA | NA | NA | 0.5* |
| red clover | *Trifolium pratense* | Multiple | 0.8<sup>15</sup> | 2* | NA | NA | NA | NA | 1.25* |
| kiwi | *Actinidia deliciosa* | Multiple | 45.4<sup>16</sup> | 1400<sup>25</sup> | NA | NA | NA | NA | 200* |
| lupine | *Lupinus spp.* | Multiple | 1.5<sup>17</sup> | 4<sup>17</sup> | NA | NA | NA | NA | 2* |
| chickpeas, lentils | *Cicer arietinum, Lens culinaris* | Multiple | 3* | 3* | NA | NA | NA | NA | 3* |
| tomato | *Solanum lycopersicum* | Saladette | 52.5<sup>12</sup> | 400* | 91.2<sup>12</sup> | NA | 133<sup>12</sup> | NA | NA |
| lowbush blueberry | *Vaccinium angustifolium* | Multiple | 5* | 65* | NA | NA | NA | NA | 15* |
| rabbiteye blueberry | *Vaccinium ashei / virgatum* | Multiple | 5* | 80* | NA | NA | NA | NA | 15* |

\* Estimate based on genus
1. Stanghellini, M.S., Ambrose, J.T., Schultheis, J.R. (1998). Seed Production in Watermelon: A Comparison between Two Commercially Available Pollinators. HortScience 33, 28–30. https://doi.org/10.21273/HORTSCI.33.1.0028
2. Rogers, S.R., Tarpy, David.R., Burrack, H.J. (2013). Multiple Criteria for Evaluating Pollinator Performance in Highbush Blueberry (Ericales: Ericaceae) Agroecosystems. Environ Entomol 42, 1201–1209. https://doi.org/10.1603/EN12303
3. Garratt, M., Fountain, M.T., McKerchar, M., Webber, S.M. (2016). Valuing insect pollinators for UK apple production. https://doi.org/10.17864/1947.50
4. Witter, S., Nunes-Silva, P., Lisboa, B.B., Tirelli, F.P., Sattler, A., Both Hilgert-Moreira, S., Blochtein, B. (2015). Stingless Bees as Alternative Pollinators of Canola. J Econ Entomol 108, 880–886. https://doi.org/10.1093/jee/tov096
5. Kandori, I., Shimaoka, R., Tsukamoto, T., Kamiya, K., Yokoi, T. (2024). Multiyear study of pollinator efficiency and importance of a wide array of pollinators in a field-cultivated strawberry plot. PLOS ONE 19, e0297130. https://doi.org/10.1371/journal.pone.0297130
6. Lundin, O. (2023). Partitioning pollination services to faba bean (*Vicia faba* L.) between managed honeybees and wild bees. Basic and Applied Ecology 71, 9–17. https://doi.org/10.1016/j.baae.2023.05.006
7. Dymond, K., Celis-Diez, J.L., Potts, S.G., Howlett, B.G., Willcox, B.K., Garratt, M.P.D. (2021). The role of insect pollinators in avocado production: A global review. Journal of Applied Entomology 145, 369–383. https://doi.org/10.1111/jen.12869
8. Holzschuh, A., Dudenhöffer, J.-H., Tscharntke, T. (2012). Landscapes with wild bee habitats enhance pollination, fruit set and yield of sweet cherry. Biological Conservation 153, 101–107. https://doi.org/10.1016/j.biocon.2012.04.032
9. Scheper, J., Badenhausser, I., Kantelhardt, J., Kirchweger, S., Bartomeus, I., Bretagnolle, V., Clough, Y., Gross, N., Raemakers, I., Vilà, M., Zaragoza-Trello, C., Kleijn, D. (2023). Biodiversity and pollination benefits trade off against profit in an intensive farming system. Proceedings of the National Academy of Sciences 120, e2212124120. https://doi.org/10.1073/pnas.2212124120
10. Klein, A.-M., Hendrix, S.D., Clough, Y., Scofield, A., Kremen, C. (2015). Interacting effects of pollination, water and nutrients on fruit tree performance. Plant Biology 17, 201–208. https://doi.org/10.1111/plb.12180
11. Artz, D.R., Nault, B.A. (2011). Performance of Apis mellifera, Bombus impatiens, and Peponapis pruinosa (Hymenoptera: Apidae) as Pollinators of Pumpkin. J Econ Entomol 104, 1153–1161. https://doi.org/10.1603/EC10431
12. Macias-Macias, O., Chuc, J., Ancona-Xiu, P., Cauich, O., Quezada-Euán, J.J.G. (2009). Contribution of native bees and Africanized honey bees (Hymenoptera:Apoidea) to Solanaceae crop pollination in tropical México. Journal of Applied Entomology 133, 456–465. https://doi.org/10.1111/j.1439-0418.2009.01399.x
13. Andrikopoulos, C.J., Cane, J.H. (2018). Comparative Pollination Efficacies of Five Bee Species on Raspberry. J Econ Entomol 111, 2513–2519. https://doi.org/10.1093/jee/toy226
14. Kasajima, S., Katagiri, C., Morishita, T., Suzuki, T., Mukasa, Y. (2017). Growth and yield of self-compatible and hybrid common buckwheat lines pollinated with and without flies. Plant Production Science 20, 384–388. https://doi.org/10.1080/1343943X.2017.1375380
15. Jing, S., Kryger, P., Markussen, B., Boelt, B. (2021). Pollination and Plant Reproductive Success of Two Ploidy Levels in Red Clover (Trifolium pratense L.). Front. Plant Sci. 12. https://doi.org/10.3389/fpls.2021.720069
16. Abbate, A.P., Campbell, J.W., Vinson, E.L., Williams, G.R. (2021). The Pollination and Fruit Quality of Two Kiwifruit Cultivars (Actinidia chinensis var. chinensis ‘AU Golden Sunshine’ and ‘AU Gulf Coast Gold’) (Ericales: Actinidiaceae) Grown in the Southeastern United States. J Econ Entomol 114, 1234–1241. https://doi.org/10.1093/jee/toab075
17. Williams, I.H., Martin, A.P., Ferguson, A.W., Clark, S.J. (1990). Effect of pollination on flower, pod and seed production in white lupin (Lupinus albus). The Journal of Agricultural Science 115, 67–73. https://doi.org/10.1017/S0021859600073937
18. Wijesinghe, S.A.E.C., Evans, L.J., Kirkland, L., Rader, R. (2020). A global review of watermelon pollination biology and ecology: The increasing importance of seedless cultivars. Scientia Horticulturae 271, 109493. https://doi.org/10.1016/j.scienta.2020.109493
19. Dogterom, M.H., Winston, M.L., Mukai, A. (2000). Effect of pollen load size and source (self, outcross) on seed and fruit production in highbush blueberry cv. ‘Bluecrop’ (VACCINIUM CORYMBOSUM; Ericaceae). American Journal of Botany 87, 1584–1591. https://doi.org/10.2307/2656734
20. Garratt, M.P.D., Bishop, J., Degani, E., Potts, S.G., Shaw, R.F., Shi, A., Roy, S. (2018). Insect pollination as an agronomic input: Strategies for oilseed rape production. Journal of Applied Ecology 55, 2834–2842. https://doi.org/10.1111/1365-2664.13153
21. Cao, Z., He, G., Mu, S., Qu, H. (2023). Effects of Bee Density and Hive Distribution on Pollination Efficiency for Greenhouse Strawberries: A Simulation Study. Agronomy 13, 731. https://doi.org/10.3390/agronomy13030731
22. Ashokkumar, G., Sundar, S.T.B., Savitha, B.K., Rameshkumar, D., Balasubramanian, P., Sampathkumar, T., Sathiyaraj, D. (2023). Genetic Variability and Diversity for Yield and Yield Attributing Traits in M3 and M4 Generations of Butter Bean (Phaseolus lunatus L.). Legume Research 46, 1308–1313.
23. Quero-García, J., Lezzoni, A., Pulawska, J., Lang, G. (2017). Cherries: botany, production and uses. CABI.
24. Goodwin, R.M., Cox, H.M., Taylor, M.A., Evans, L.J., McBrydie, H.M. (2011). Number of honey bee visits required to fully pollinate white clover (Trifolium repens) seed crops in Canterbury, New Zealand. New Zealand Journal of Crop and Horticultural Science 39, 7–19. https://doi.org/10.1080/01140671.2010.520164
25. Ferguson, A.R. (2013). Chapter Two - Kiwifruit: The Wild and the Cultivated Plants, in: Boland, M., Moughan, P.J. (Eds.), Nutritional Benefits of Kiwifruit, Advances in Food and Nutrition Research. Academic Press, pp. 15–32. https://doi.org/10.1016/B978-0-12-394294-4.00002-X
26. Eeraerts, M., Vanderhaegen, R., Smagghe, G., Meeus, I. (2020). Pollination efficiency and foraging behaviour of honey bees and non-Apis bees to sweet cherry. Agricultural and Forest Entomology 22, 75–82. https://doi.org/10.1111/afe.12363
27. Bosch, J., Blas, M. (1994). Foraging Behaviour and Pollinating Efficiency of Osmia cornuta and Apis mellifera on Almond (Hymenoptera, Megachilidae and Apidae). Applied Entomology and Zoology 29, 1–9. https://doi.org/10.1303/aez.29.1
28. Garantonakis, N., Varikou, K., Birouraki, A., Edwards, M., Kalliakaki, V., Andrinopoulos, F. (2016). Comparing the pollination services of honey bees and wild bees in a watermelon field. Scientia Horticulturae 204, 138–144. https://doi.org/10.1016/j.scienta.2016.04.006
29. Kingha, B.M.T., Fohouo, F.-N.T., Brückner, A.N. and D. (2012). Foraging and pollination activities of Xylocopa olivacea (Hymenoptera, Apidae) on Phaseolus vulgaris (Fabaceae) flowers at Dang (Ngaoundere-Cameroon). JAERD 4, 330–339. https://doi.org/10.5897/JAERD11.151

**Table S2:**
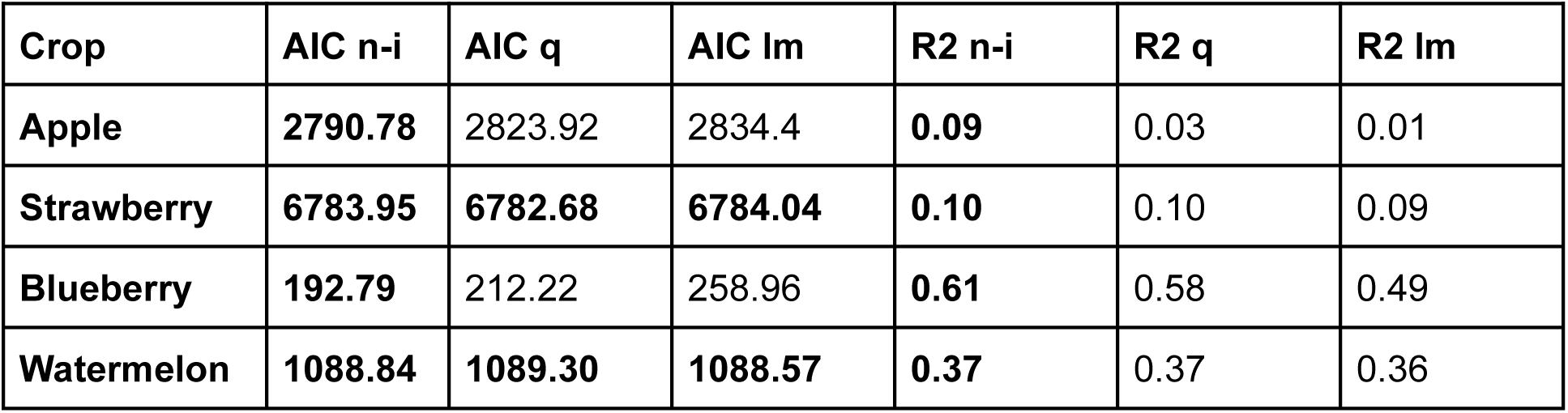
AIC and R^2^ Comparison of different fits to empirical data. Non-inflected (n-i), quadratic (q), linear model (lm). In bold, the best model according to AIC (for Delta AIC < 2, all models are selected).

| **Crop** | **AIC n-i** | **AIC q** | **AIC lm** | **R2 n-i** | **R2 q** | **R2 lm** |
|---|---|---|---|---|---|---|
| **Apple** | **2790.78** | 2823.92 | 2834.4 | **0.09** | 0.03 | 0.01 |
| **Strawberry** | **6783.95** | **6782.68** | **6784.04** | **0.10** | 0.10 | 0.09 |
| **Blueberry** | **192.79** | 212.22 | 258.96 | **0.61** | 0.58 | 0.49 |
| **Watermelon** | **1088.84** | **1089.30** | **1088.57** | **0.37** | 0.37 | 0.36 |

**Fig. S1:**
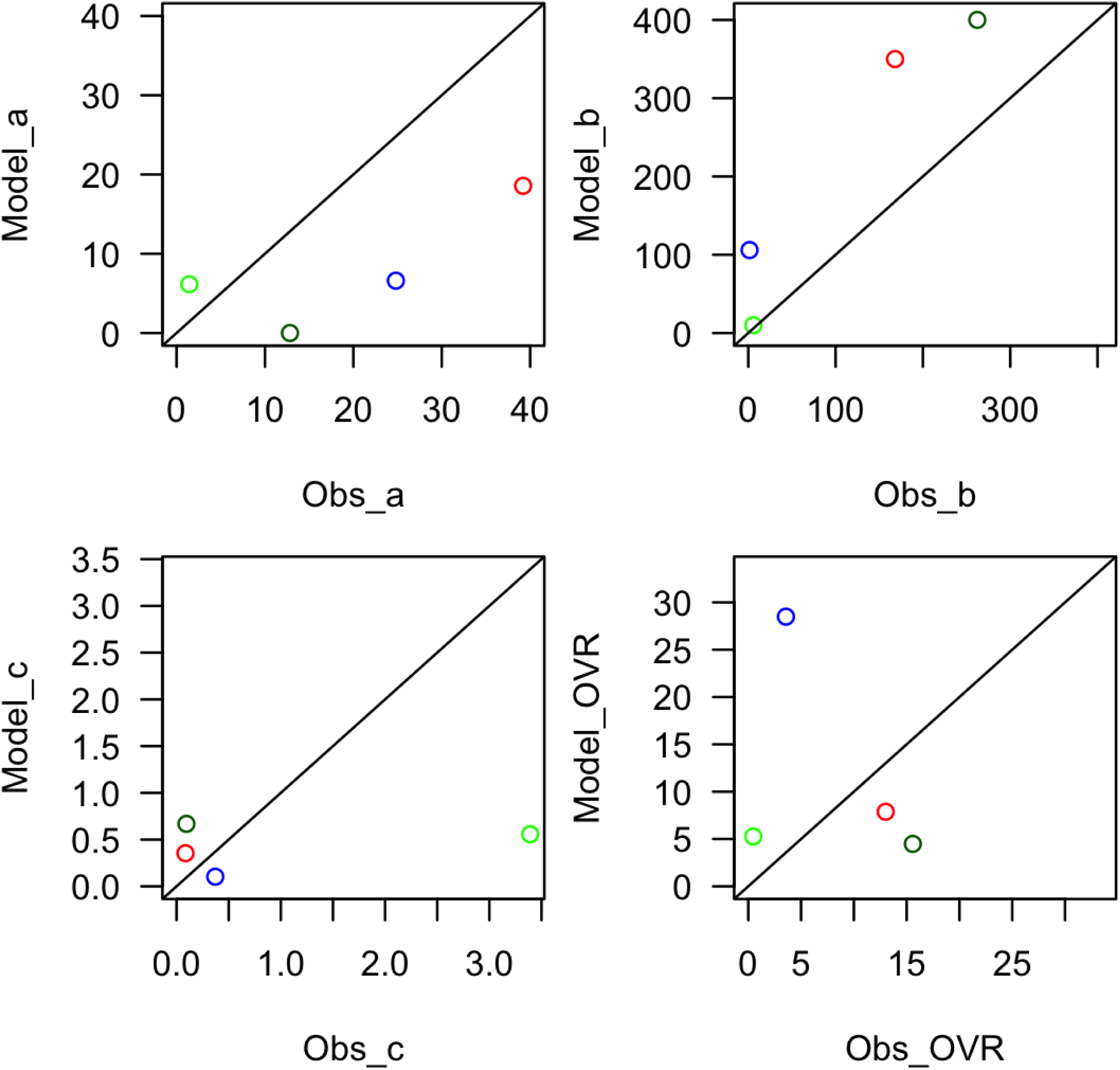
Comparison of observed values fitting empirical data (See Fig. 2) against mechanistic prediction using independent data from other studies. The bottom line is that different cultivars might vary a lot in raw values, and the models are sensitive to those inputs. For example, single-visit efficiency for watermelon is ∼200 seeds, while the empirically observed averages ∼70 seeds, explaining the large discrepancy.

**Fig S2:**
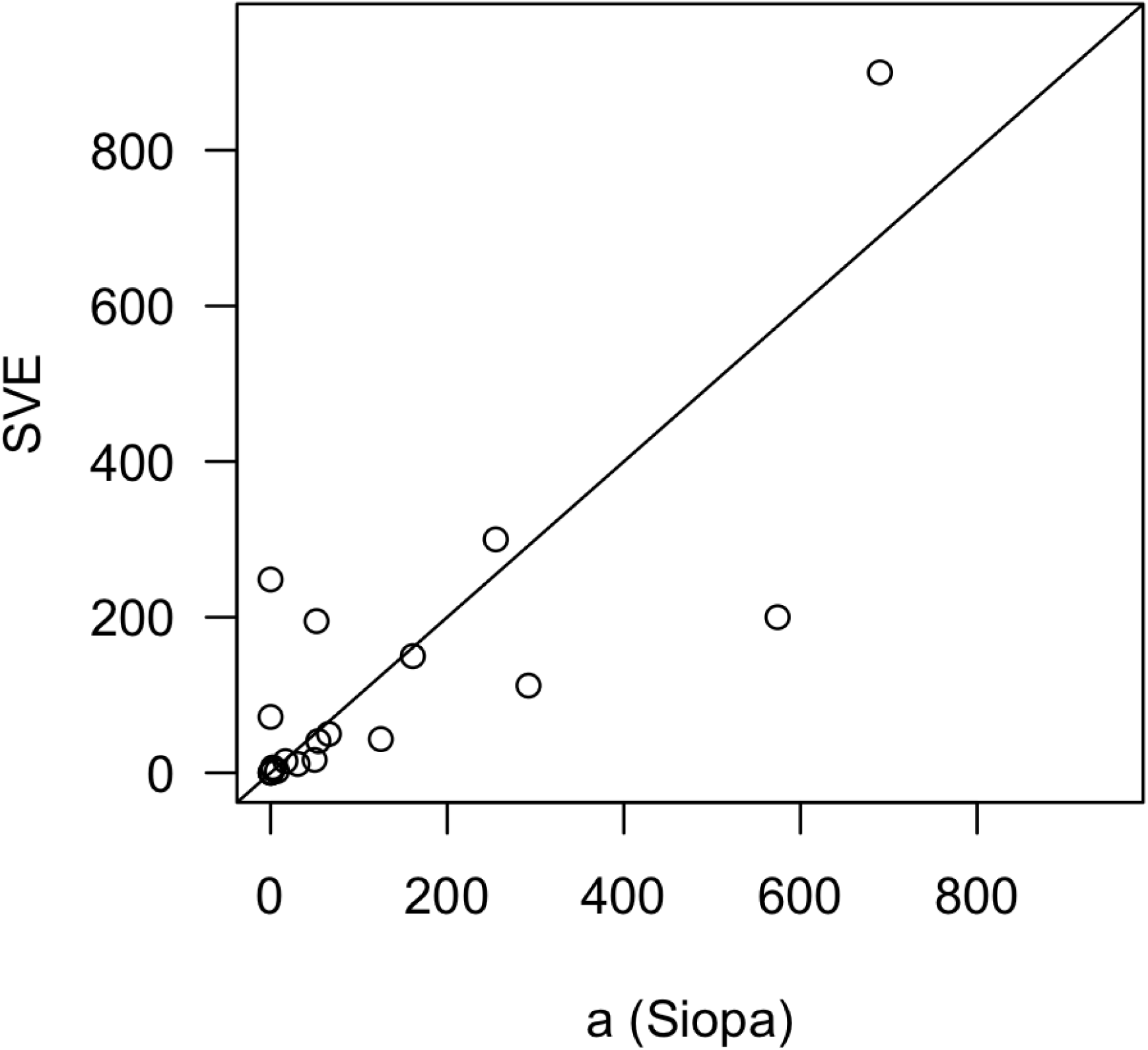
The number of seeds produced with no pollination (*a*) according to Siopa et al. (REF) is higher than the seed set produced by a single visit of a pollinator in 11 out of 26 studies (dots below the 1:1 line). This is due to the large variability between studies in reported selfing rates. This cautions against using averages across crops, as in Siopa, in combination with single-study data from a single cultivar.

